# Commercial vaccines do not confer protection against two genetic strains of *Piscirickettsia salmonis*, LF-89-like and EM-90-like, in Atlantic salmon

**DOI:** 10.1101/2021.01.07.424493

**Authors:** Carolina Figueroa, Débora Torrealba, Byron Morales-Lange, Luis Mercado, Brian Dixon, Pablo Conejeros, Gabriela Silva, Carlos Soto, José A. Gallardo

## Abstract

In Atlantic salmon, vaccines have failed to control and prevent Piscirickettsiosis, for reasons that remain elusive. In this study, we report the efficacy of a commercial vaccine developed with the *Piscirickettsia salmonis* isolate AL100005 against other two isolates which are considered highly and ubiquitously prevalent in Chile: LF-89-like and EM-90-like. Two cohabitation trials were performed to mimic real-life conditions and vaccine performance: 1) post smolt fish were challenged with a single infection of LF-89-like, 2) adults were coinfected with EM-90-like and a low coinfection of sea lice. In the first trial, the vaccine delayed smolt mortalities by two days; however, unvaccinated and vaccinated fish did not show significant differences in survival (unvaccinated: 60.3%, vaccinated: 56.7%; p = 0.28). In the second trial, mortality started three days later for vaccinated fish than unvaccinated fish. However, unvaccinated and vaccinated fish did not show significant differences in survival (unvaccinated: 64.6%, vaccinated: 60.2%, p= 0.58). Thus, we found no evidence that the evaluated vaccines confer effective protection against of LF-89-like or EM-90-like with estimated relative survival proportions (RPSs) of −9% and −12%, respectively. More studies are necessary to evaluate whether pathogen heterogeneity is a key determinant of the vaccine efficacy against *P. salmonis.*

## 1 Introduction

*Piscirickettsia salmonis* is a major concern for the Chilean salmon industry, causing economic losses of USD 700 million per year (Maisey et al., 2017; Rozas and Enriquez, 2014). Piscirickettsiosis is an exceptionally contagious disease, with a high prevalence in clustered regions in Chile, causing mortalities of over 50% of production (Jakob et al., 2014; Leal and Woywood, 2007). While Chile, the second-largest global producer of salmon, is by far the most affected country by this disease, it also affects the other main salmon producing countries, namely Norway, Canada and Scotland (Brocklebank et al., 1993; Grant et al., 1996; Olsen et al., 1997; SERNAPESCA, 2017).

Vaccination has been widely used as a control strategy to prevent Piscirickettsiosis (Happold et al., 2020), but unfortunately, all vaccines developed in the last 20 years have failed to protect Atlantic salmon against *P. salmonis* (Maisey et al., 2017). Some intrinsic and extrinsic factors that may explain why commercial vaccines present reduced protection against *P. salmonis* are 1) coinfection with sea lice, which were able to override the protective effects of vaccines (Figueroa et al., 2017; Figueroa et al., 2020 accepted); 2) host genetic variation, partially protecting some hosts while leaving others unprotected (Figueroa et al., 2017; Figueroa et al., 2020 accepted); and 3) ineffectiveness in stimulating cellular immunity, which is a key element to protecting against *P. salmonis* because this bacteria can survive inside the host cells. Likewise, other underlying causes may lead to low vaccine efficacy, such as the pathogen’s genetic variation.

Since Piscirickettsia outbreaks in Chile are caused by a minimum of two different genetic strains, it has been suggested that this heterogeneity should be considered in vaccine development (Nourdin-Galindo et al., 2017; Otterlei et al., 2016). The reported efficacy of a commercial vaccine would not hold in the field when testing against bacterial strains with low virulence and/or a reduced prevalence in the field. In Chile, two strains—called LF-89-like and EM-90-like—are considered highly and ubiquitously prevalent (Saavedra et al., 2017). These strains show distinct laboratory growth conditions (Saavedra et al., 2017) and major differences in virulence-associated secretion systems and transcriptional units (Millar et al., 2018), resulting in different infective levels (Bohle et al., 2014). For example, it has been shown that EM-90-like isolates are more aggressive than the LF-89-like isolates, inducing higher cumulative mortalities (EM-90 = 95%; LF89 = 82%) and a shorter time to death (EM-90 = 42 days; LF89 = 46 days) in non-vaccinated post-smolt when evaluated by a cohabitation challenge (Rozas-Serri et al., 2017; Rozas-Serri et al., 2018). Contrary to the hypothesis of heterogeneity, an experimental vaccine developed with an isolate of EM-90 failed to protect against the same isolate (Cardella and Eimers, 1990; Meza et al., 2019).

In this study, we tested the efficacy of a commercial vaccine developed with the *P. salmonis* isolate AL100005 against the two most prevalent Chilean strains of Piscirickettsia, LF-89-like and EM-90-like. The challenges were carried out with Atlantic salmon (*Salmo salar*) and by cohabitation with fish that successfully adapted to saline conditions in order to best imitate the natural conditions of bacterial infection. In the first trial, LF-89-like was evaluated with post-smolt fish with a single infection of *P. salmonis,* while in the second trial, EM-90-like was evaluated with adult fish in a challenge that included a very low coinfection with the sea louse *C. rogercresseyi,* again to better emulate field conditions.

## 2 Materials and methods

### Ethics Statement

This work was carried out under the guidance for the care and use of experimental animals of the Canadian Council on Animal Care. The protocol was approved by the Bioethics Committee of the Pontificia Universidad Católica de Valparaíso and the Comisión Nacional de Investigación Científica y Tecnológica de Chile (FONDECYT N° 1140772). Animals were fed daily *ad libitum* with a commercial diet. To reduce stress during handling, vaccination was performed on fish that were sedated with AQUI-S (50% Isoeugenol, 17 mL/100 L water). Euthanasia was performed using an overdose of anesthesia.

### Commercial vaccine

The commercial vaccine, hereafter “the vaccine”, used in this study was a pentavalent vaccine with antigens against *P. salmonis, Vibrio ordalii, Aeromonas salmonicida*, IPNV (Infectious Pancreatic Necrosis Virus) and ISAV (Infectious Salmon Anemia Virus). This vaccine is the most used (3821/8884 or 43 % of vaccination events in the freshwater phase of production) by Chilean farmers, as was reported by (Happold et al., 2020) The active principle of this vaccine to prevent Piscirickettsiosis is a inactivated vaccine of *P. salmonis* AL 10005 strain.

### Challenge with LF-89-like (Trial 1)

A total of 4983 individually pit-tagged smolt fish were provided in 2017 by the Salmones Camanchaca Company. Fish were transferred to the experimental station of the Neosalmon Company for the evaluation of the vaccine efficacy using a cohabitation challenge (Table 1). In total, 1223 of the fish had been previously immunized with the vaccine using the normal production schedule and were used as vaccinated fish (342 ± 55 g), 861 had been previously injected with Phosphate-Buffered Saline (PBS) and were used as unvaccinated fish (314 ± 61 g), while the remaining fish were used as Trojan shedders (152 ± 38 g).

**Table 1.**
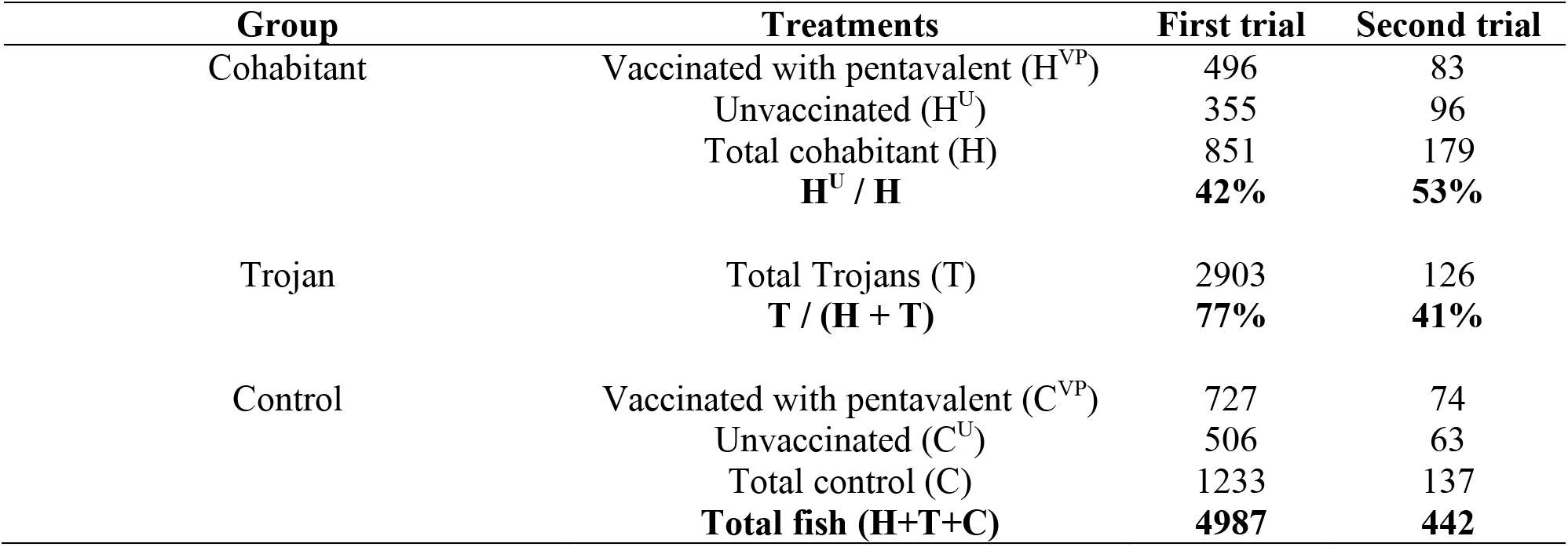
Number and proportion of Atlantic salmon used per group and treatment for the first and second trials. In the first trial, post-smolt fish were challenged with the LF-89-like isolate of *P. salmonis*, while in the second trial, adult fish were challenged with the EM-90-like isolate of *P. salmonis* and with the sea lice *C. rogercresseyi*.

| Group | Treatments | First trial | Second trial |
| --- | --- | --- | --- |
| Cohabitant | Vaccinated with pentavalent (H<sup>VP</sup>) | 496 | 83 |
|  | Unvaccinated (H<sup>U</sup>) | 355 | 96 |
|  | Total cohabitant (H) | 851 | 179 |
|  | **H<sup>U</sup> / H** | **42%** | **53%** |
| Trojan | Total Trojans (T) | 2903 | 126 |
|  | **T / (H + T)** | **77%** | **41%** |
| Control | Vaccinated with pentavalent (C<sup>VP</sup>) | 727 | 74 |
|  | Unvaccinated (C<sup>U</sup>) | 506 | 63 |
|  | Total control (C) | 1233 | 137 |
|  | **Total fish (H+T+C)** | **4987** | **442** |

Vaccinated and unvaccinated fish were distributed in four tanks: two tanks of 15 m^3^ for the cohabitation challenges and two tanks of 5 m^3^ for the control without infection. All fish were acclimatized to the experimental conditions (salinity of 32% and a temperature of 15 ± 1 °C) and tanks for at least 15 days prior to the challenge. Further, a health check by RT-PCR was performed to verify that the fish were free of viral (ISAV and IPNV) and bacterial pathogens (*Vibrio* sp., *Flavobacterium* sp., *P. salmonis,* and *Renibacterium salmoninarum*). The cohabitation tanks were challenged by adding Trojan shedders (table 1) which had been previously anesthetized with AQUI-S and injected with a median lethal dose (LD_50_) of 1×10^−2^ TCID/ml (TCID: median tissue culture infective dose) of LF-89-like isolate provided by ADL Diagnostics Company. The experiment was conducted 43 days after the *P. salmonis* injection of Trojans.

The LD_50_ used in Trojans was previously determined on 800 immunized fish with the vaccine, which were equally distributed in four treatments and two tanks of 1000 L per treatment. Treatment 1 involved injection with 1 × 10^−2^ TCID/ml, treatment 2 involved injection with 1 × 10^−3^ TCID/ml, treatment 3 involved injection with 1 × 10^−4^ TCID/ml, and treatment 4 involved injection with Phosphate-Buffered Saline (PBS). Fish were monitored daily for 30 days, and mortalities were recorded.

### Challenge with EM-90-like and coinfection with sea lice (Trial 2)

A total of 442 individually pit-tagged adult fish were provided in 2019 by the company Salmones Camanchaca and transferred to the experimental station of the Aquadvice company for the evaluation of the efficacy vaccine using a cohabitation challenge (Table 1). In total, 170 of the fish had been previously immunized with the vaccine using the normal production schedule and were used as vaccinated fish (1,274 ± 318 g), 146 had been previously injected with Phosphate-Buffered Saline (PBS) and were used as unvaccinated fish (1,260 ± 345 g), while the remaining fish were used as Trojan shedders (1,311 ± 346 g).

Vaccinated and unvaccinated fish were distributed in three tanks of 11 m^3^: two tanks for the cohabitation challenges and one tank for the control without infection. All fish were acclimatized to the experimental conditions (salinity of 32% and a temperature of 15 ± 1 °C) and tanks for at least 15 days prior to the challenge. Further, a health check by RT-PCR was performed to verify that the fish were free of viral (ISAV and IPNV) and bacterial pathogens (*Vibrio* sp., *Flavobacterium* sp., *P. salmonis,* and *Renibacterium salmoninarum*). The cohabitation tanks were challenged by adding Trojan shedders (table 1) which had been previously anesthetized with AQUI-S and injected with a median lethal dose (LD_50_) of 1 × 10^−3.5^ TCID/ml of EM-90-like isolate provided by Fraunhofer, Chile. After seven days of the Trojan fish being challenged with *P. salmonis,* all fish (cohabitant, Trojan and control) were infested with copepodids of *C. rogercresseyi*. The coinfection procedure was established based on our previous studies (Araya et al., 2012; Lhorente et al., 2014), but now a very low infection rate was applied to mimic the natural infection rates normally seen in field conditions (Bravo et al., 2010). Infections with sea lice were performed by adding 20 copepodites per fish to each control and coinfection tank. Copepodites were collected from egg-bearing females reared in the laboratory and confirmed as pathogen-free (*P. salmonis*, *R. salmoninarum,* IPNV, and ISAV) by RT-PCR diagnosis. After the addition of parasites, water flow was stopped for a period of 8 h, and tanks were covered to decrease light intensity, which favors the successful settlement of sea lice on fish (Araya et al., 2012). Parasite counting was performed a week after the infestation in a sample of nine fish per tank. The challenge lasted 60 days after the Trojans’ infection with *P. salmonis*.

The LD_50_ used in Trojans was previously determined on 330 immunized fish with the vaccine, which were equally distributed in five treatments and two tanks of 720 L per treatment. Treatment 1 involved injection with 1 × 10^−1.5^ TCID/ml, treatment 2 involved injection with 1 × 10^−2.5^ TCID/ml, treatment 3 involved injection with 1×10^−3.5^ TCID/ml, treatment 4 involved injection with 1 × 10^−4.5^ TCID/ml, and treatment 5 involved injection with Phosphate-Buffered Saline (PBS). Fish were monitored daily for 30 days, and mortalities were recorded.

### Necropsy analysis

Macroscopic lesions from 10 controls and cohabitant fish in each trial were analyzed. Two different veterinarians who were blinded to the treatments studied fresh samples from trials 1 or 2. In the challenge with LF-89-like, macroscopic lesions were evaluated at 21 days post-infection in the liver where vacuolar degeneration, hepatitis and hepatocyte atrophy were described according to their presence or absence. Further, 47 vaccinated and unvaccinated fish from cohabitation and control tanks were analyzed by immunohistochemistry to detect the presence or absence of *P. salmonis* in the liver at 21 days after challenges and at the end of the experiment. In the challenge with EM-90-like, clinical signs were evaluated at the end of the challenges; the analysis included presence or absence of nodules in the liver, congestive liver, and hepatomegaly.

### ELISA

An indirect Enzyme-Linked Immunosorbent Assay (ELISA) was performed in serum samples only from the first trial—the fish challenged with LF-89-like isolate. Secretion levels of total immunoglobulin (Igs), antigen-specific immunoglubulins against *P. salmonis* (spIgs), tumor necrosis factor-alpha (TNFα) and interferon-gamma (IFNγ) were measured following the protocol of Morales-Lange *et al*. (Morales-Lange et al., 2018). Briefly, the total protein concentration of each sample was first determined by the BCA (Bicinchoninic acid) method (Pierce, Thermo Fisher, Waltham, USA) according to the supplier’s instructions. Then, each sample was diluted in carbonate buffer (60 mM NaHCO_3_, pH 9.6), seeded in duplicate at 50 ng μL^−1^ (100 μL) in a Maxisorp plate (Nunc, Thermo Fisher Scientific, Waltham, USA) and incubated overnight at 4 °C. After that, the plates were blocked with 200 μL per well of 1% Bovine Serum Albumin (BSA) for 2 h at 37 °C, and later the primary antibodies (Supplementary table and figure 1) were incubated for 90 min at 37 °C. Next, a secondary antibody—HRP (Thermo Fisher)—was incubated for 60 min at 37 °C in a 1:7000 dilution. Finally, 100μL per well of chromagen substrate 3,30,5,50-tetrame thylbenzidine (TMB) single solution (Invitrogen, California, USA) was added and incubated for 30 min at room temperature. The reaction was stopped with 50 μL of 1 N sulfuric acid and read at 450 nm on a VERSAmax microplate reader (Molecular Device, California, USA). For the detection of spIg, 50 ng μL^−1^ of total protein extract from *P. salmonis* (Carril et al., 2017) were seeded per well in a Maxisorp plate (diluted in 100 μL of carbonate buffer) and incubated overnight at 4 °C. After blocking with 1% BSA (200 μL per well), each fish serum sample was incubated in duplicate at a total Igs concentration of 50 ng μL^−1^ for 90 min at 37 °C. After that, the ELISA protocol described above was followed.

**Figure 1.**
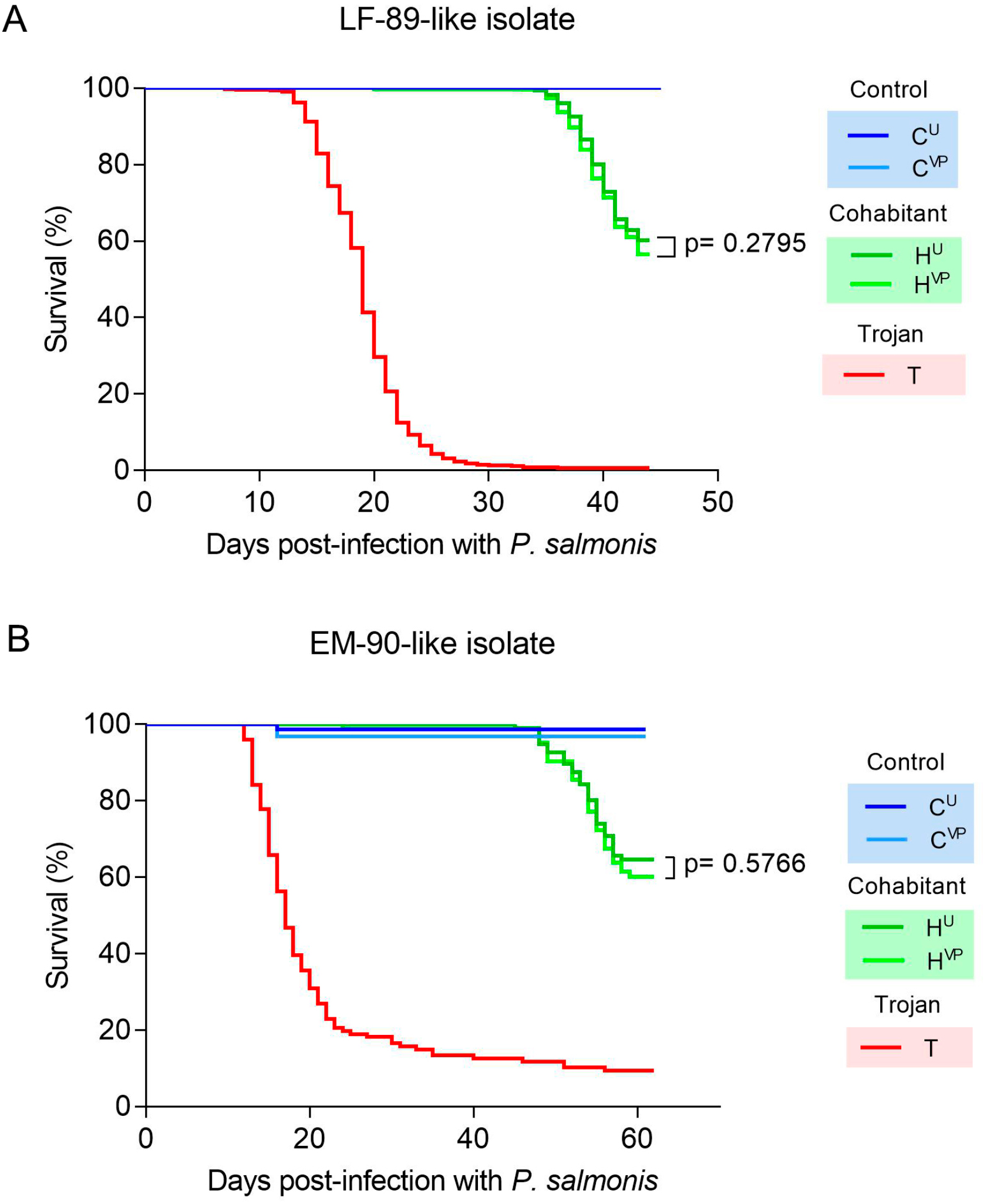
Survival curves. **(A)** Single infection of Atlantic salmon post-smolt with the *P. salmonis* LF-89-like strain. **(B)** Coinfection of Atlantic salmon adults with the *P. salmonis* EM-90-like strain and the sea louse *C. rogercresseyi*. Significant differences were obtained from the Log-rank test. Abbreviations: C^U^: control unvaccinated; C^VP^: control vaccinated with pentavalent; H^U^: cohabitant unvaccinated; H^VP^: cohabitant vaccinated with pentavalent; T: trojan.

### Statistical analysis

The mortality was registered in all individuals, and data were represented using Kaplan–Meier survival curves (Kaplan and Meier, 1958). The protection elicited by vaccines was determined by comparing the percentage of survival of vaccinated and unvaccinated groups using a Log-rank test. Further, the relative proportion survival (RPS) was calculated as

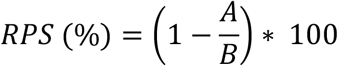

where A and B are the mortalities at the end of challenges in vaccinated and unvaccinated fish, respectively.

Additionally, differences in the clinical signs of *P. salmonis* infection between different treatments were analyzed using a non-parametric Chi-square test. Finally, significant differences in ELISA tests were compared using the Student’s two-tailed t-test, *p* < 0.05. All statistical analyses were performed using R Core Team (RStudio, Vienna, Austria). Graphs were designed with GraphPad Prism 8.0 software (GraphPad Software, CA, USA).

## 3 Results

### Vaccine efficacy against LF-89-like isolate

As we expected, the cohabitation challenge with the LF-89-like isolate of *P. salmonis* resulted in high mortality in the cohabiting fish and no mortality in the non-infected control fish. However, we found no evidence that the evaluated vaccine generated an effective protection against this strain. The vaccine delayed mortalities by two days (H^U^: 34 dpi and H^VP^: 36 dpi), but unvaccinated fish and those vaccinated showed similar survival during and at the end of the challenges (H^VP^: 56.7% and H^U^: 60.3%, Figure 1A). The survival test did not reveal significant differences between vaccinated and unvaccinated treatments (p = 0.28).

Dead fish and large numbers of vaccinated and unvaccinated live fish at the end of the challenge showed multiple hemorrhagic ulcers on the skin typical of a severe *P. salmonis* infection. Infection with *P. salmonis* was also evident in both vaccinated and unvaccinated fish in the liver at the end of the challenge, but not at 21 days after infection (Figure 2). On the other hand, vaccination increased the presence of hepatocyte atrophy in comparison with unvaccinated fish in the control treatment at 21 days post-infection (Table 2). A similar trend was observed in the cohabitant fish, but without significant differences (Table 2). The health status of fish was evaluated again against most common salmon diseases, revealing the appearance of secondary infections of *Piscine orthoreovirus* (Supplementary Figure 3) and *Tenacibaculum dicentrarchi* in some fish.

**Table 2.**
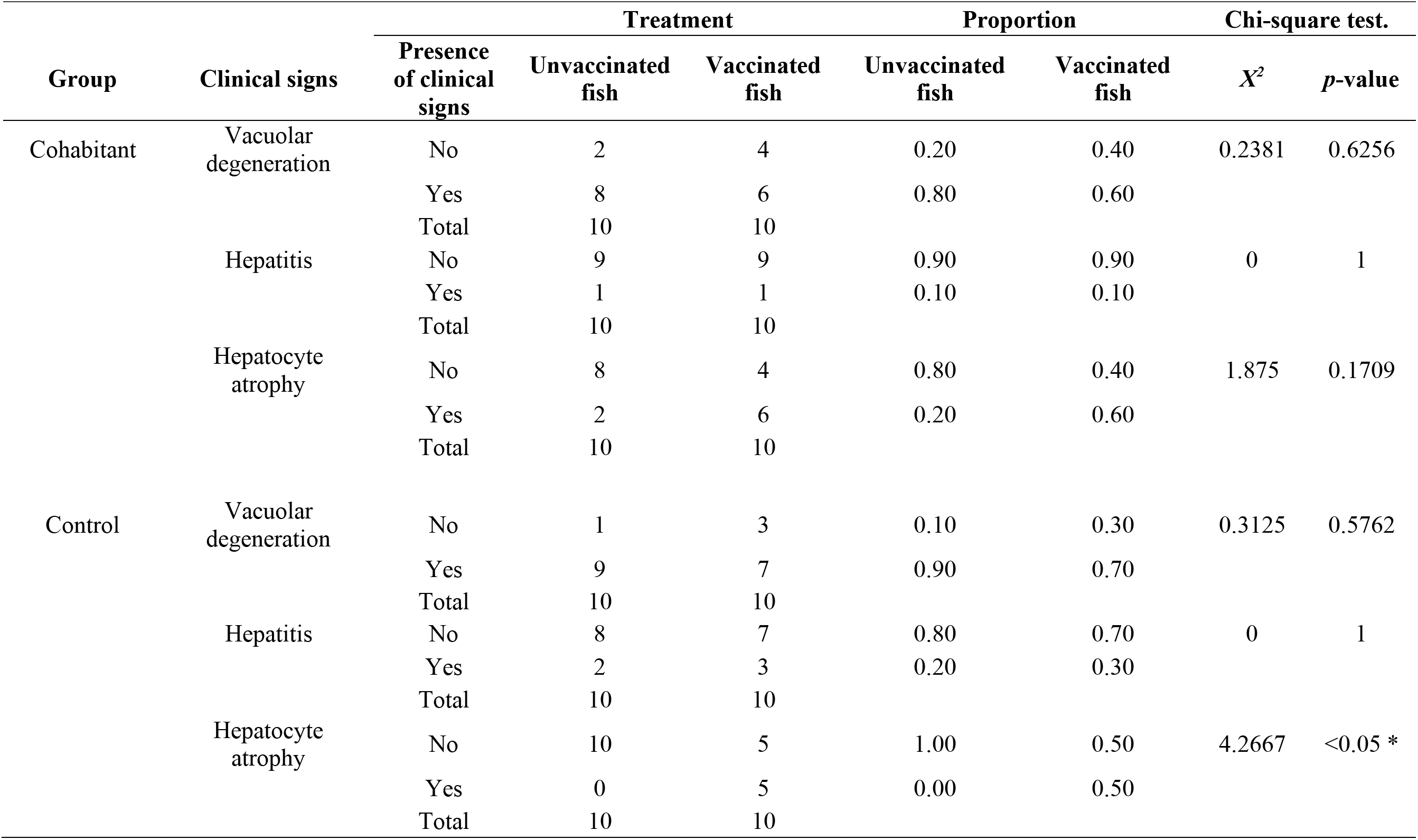
Clinical signs in Atlantic salmon challenged with the LF-89-like isolate of *P. salmonis* at day 21 post-infection in cohabitant and control groups. Differences between vaccinated and unvaccinated fish were evaluated with a Chi-squared statistical test.

**Figure 2.**
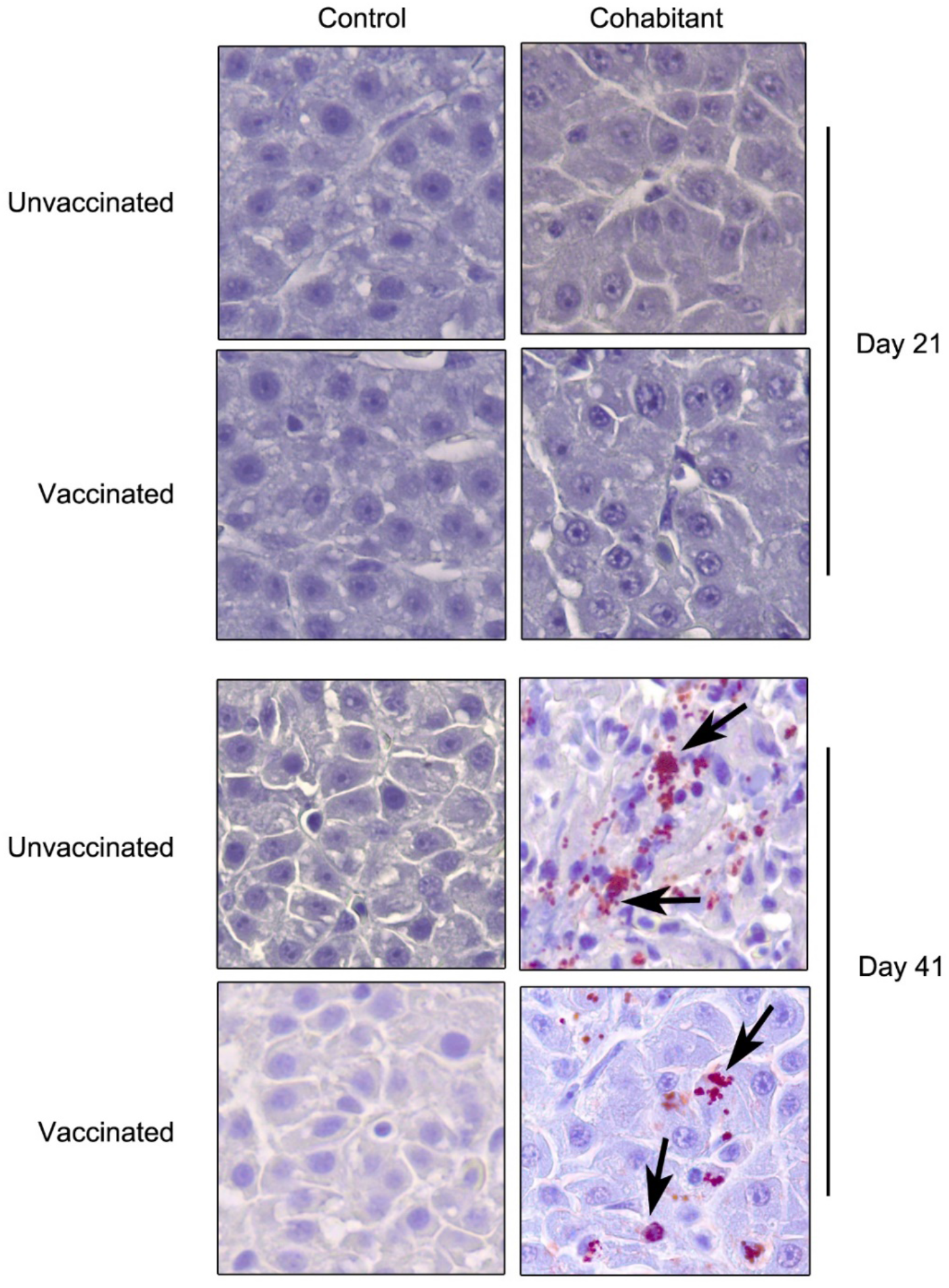
Presence of LF-89-like isolate of *P. salmonis* (black arrows) in liver samples of Atlantic salmon. Piscirickettsiosis was detected in 11 out of 47 fish analyzed by immunohistochemistry — magnification 63X.

The ELISA results in serum samples of *S. salar* showed a significant increase of total Igs at 21 days post-infection (Figure 3A) in the control group of vaccinated fish (C^VP^). However, at the same sampling time, cohabiting fish (unvaccinated and vaccinated) showed a decrease in total Igs levels. This trend was reversed at 41 dpi, since both groups (H^U^ and H^VP^) significantly increased their levels of total Igs. On the other hand, regarding specific immunoglobulins against *P. salmonis* (Figure 3B), an increase was detected in C^VP^ before the challenge with *P. salmonis*. Nevertheless, after 41 days post-infection, H^VP^ group showed less availability of spIgs against *P. salmonis* than the other groups. Finally, the evaluation of TNFα and IFNγ secretion did not show significant changes between treatments (Figure 3C-D).

**Figure 3.**
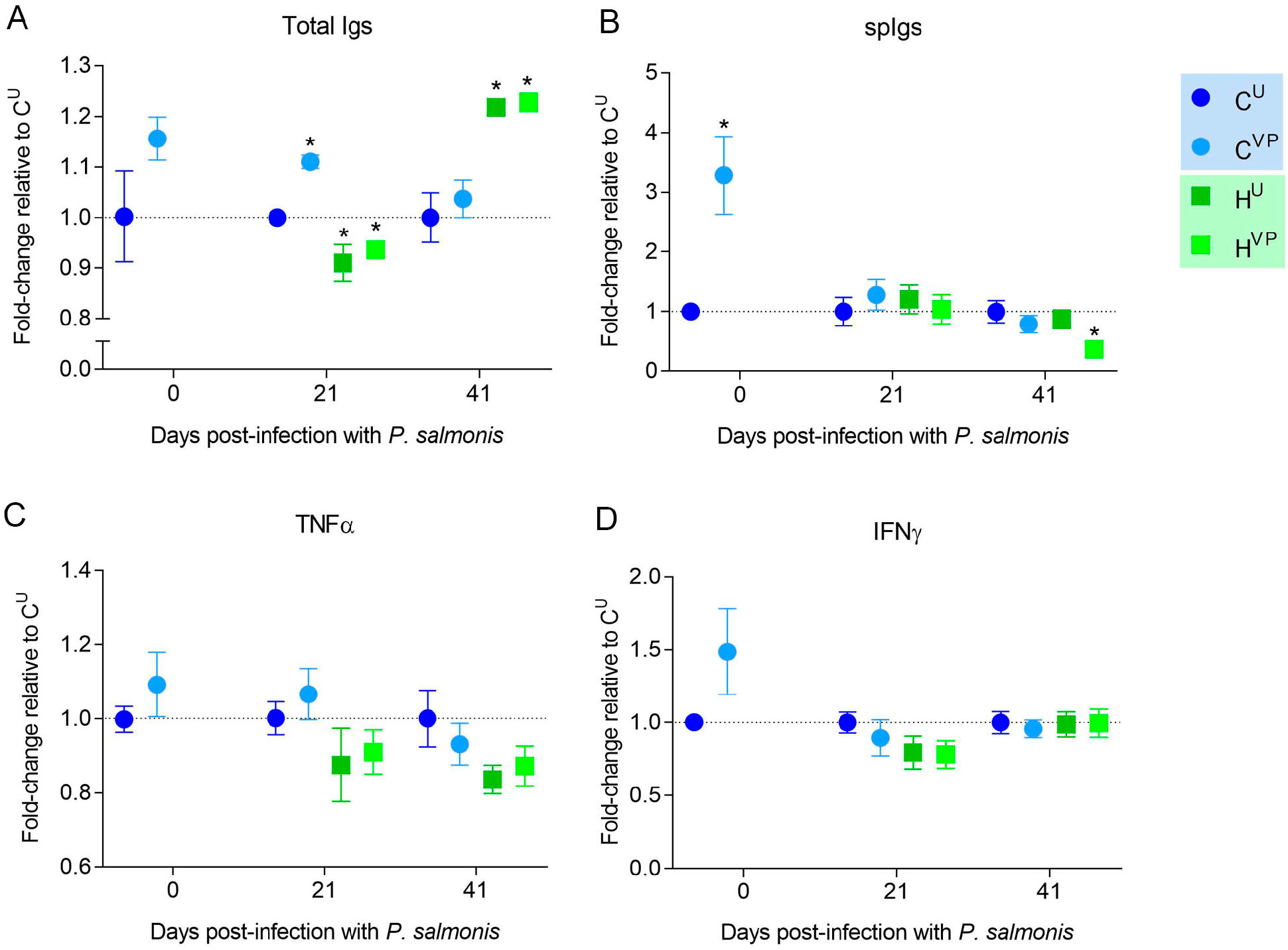
Secretion of total Igs **(A)**, antigen specific Igs **(B)**, tumor necrosis factor alpha (TNFα) **(C)** and interferon gamma (IFNγ) **(D)** in serum samples from Atlantic salmon measured by ELISA after a challenge with *P. salmonis* in the first trial (single infection of the LF-89-like isolate). Data represent the mean ± SEM (n = 10). Significant differences compared to C^U^ by Student t-test two-tailed (*p* <0.05). Abbreviations: C^U^: control unvaccinated; C^VP^: control vaccinated with pentavalent; H^U^: cohabitant unvaccinated; H^VP^: cohabitant vaccinated with pentavalent.

### Vaccine efficacy against EM-90-like isolate with low sea lice coinfection

In the second trial, adult fish were coinfected with sea lice to mimic natural conditions in the field. Seven days after sea lice infestation, the prevalence of sea lice was 100% in treatment and control tanks, with no significant differences in the abundance of the parasites between tanks (Tank 1 = 10.4 ± 4.0; Tank 2 = 11.7 ± 3.0; control tank = 9.7 ± 6.6). In cohabitant fish, the vaccine was not able to protect Atlantic salmon against the EM-90-like strain (H^VP^: 60.2% and H^U^: 64.6%; Figure 1B; p = 0.58) during a very low-level sea lice infection. However, a small protective effect was observed; for example, steady mortality started three days later for vaccinated fish compared with unvaccinated fish (H^VP^: 48 dpi and H^U^: 45 dpi). The control tank infected only with sea lice presented very low mortalities, with one fish dead of the unvaccinated fish (C^U^) and two dead of the vaccinated fish (C^VP^).

Vaccinated and unvaccinated dead fish showed hemorrhagic ulcers on the skin typical of a severe *P. salmonis* infection. Further, when we compared cohabitant and control fish at the end of the challenges, infection with *P. salmonis* was evident in the cohabitant fish in terms of the three evaluated clinical signs: nodules in liver, congestive liver and hepatomegaly (Table 3). However, we did not find differences between vaccinated and unvaccinated fish in cohabitant fish (Table 3). For instance, in the cohabitant treatment, nine unvaccinated fish presented a congestive liver, compared to 10 vaccinated fish with that condition. Similar patterns were found for nodules in the liver and hepatomegaly.

**Table 3.** Clinical signs in Atlantic salmon challenged with the EM-90-like isolate of *P. salmonis* and infestation with *C. rogercresseyi* at day 47–51 post-infection in cohabitant and control groups. Differences between vaccinated and unvaccinated fish were evaluated with a Chi-squared statistical test.

| Group | Clinical signs | Presence of clinical signs | Treatment |  | Proportion |  | Chi-square test. |  |
| --- | --- | --- | --- | --- | --- | --- | --- | --- |
| | | | Unvaccinated fish | Vaccinated fish | Unvaccinated fish | Vaccinated fish | $X^2$ | $p$-value |
| Cohabitant | Nodules in liver | No | 0 | 0 | 0 | 0 | 0 | 1 |
|  |  | Yes | 10 | 10 | 1 | 1 |  |  |
|  |  | Total | 10 | 10 |  |  |  |  |
|  | Congestive liver | No | 1 | 0 | 0.1 | 0 | 0.0023 | 0.961 |
|  |  | Yes | 9 | 10 | 0.9 | 1 |  |  |
|  |  | Total | 10 | 10 |  |  |  |  |
|  | Hepatomegaly | No | 0 | 0 | 0 | 0 | 0 | 1 |
|  |  | Yes | 10 | 10 | 1 | 1 |  |  |
|  |  | Total | 10 | 10 |  |  |  |  |
| Control | Nodules in liver | No | 10 | 10 | 1 | 1 | 0 | 1 |
|  |  | Yes | 0 | 0 | 0 | 0 |  |  |
|  |  | Total | 10 | 10 |  |  |  |  |
|  | Congestive liver | No | 10 | 9 | 1 | 0.9 | 0 | 1 |
|  |  | Yes | 0 | 1 | 0 | 0.1 |  |  |
|  |  | Total | 10 | 10 |  |  |  |  |
|  | Hepatomegaly | No | 10 | 9 | 1 | 0.9 | 0 | 1 |
|  |  | Yes | 0 | 1 | 0 | 0.1 |  |  |
|  |  | Total | 10 | 10 |  |  |  |  |

## 4 Discussion

Vaccination is one of the most relevant strategies to prevent and control diseases in aquaculture (Assefa and Abunna, 2018). However, vaccines have failed to control and prevent Piscirickettsiosis, for reasons that remain elusive (Adams, 2019; Alvarez et al., 2016; Cabello and Godfrey, 2019; Maisey et al., 2017). This manuscript evaluated whether the heterogeneity of *P. salmonis* could explain the low vaccine efficacy of a commercial vaccine whose active principle is a bacterin developed using the *P. salmonis* AL 10005 strain. To do that, we evaluated the vaccine efficacy using the two most prevalent and ubiquitous isolates of *P. salmonis* in Chile. Challenges were designed to mimic the natural condition of infection; thus, LF-89-like was evaluated with post-smolt fish in a single infection of *P. salmonis,* and EM-90-like was evaluated with adult fish in a challenge that included a very low coinfection pressure with the sea louse *C. rogercresseyi*. Thus, in this study, we found no evidence that the vaccine developed with the *P. salmonis* AL 10005 strain confers protection against LF-89-like or EM-90-like in Atlantic salmon.

The absent or low level of protection provided by vaccines against Piscirickettsia could be related to the selection of an incorrect model for the evaluation of protection in vaccination trials, which may lead to the overestimation of the real protective value of vaccines in the field. For example, the route of infection has been proposed as a relevant factor for defining the performance of a vaccine. Here, we selected a cohabitation model of challenges, because cohabitation challenges best mimic the natural infection route (Nordmo, 1997). On the other hand, several studies evaluating vaccine efficacy against *P. salmonis* have been performed by intraperitoneal injection (Kuzyk et al., 2001; Salonius et al., 2005; Tobar et al., 2011; Wilhelm et al., 2006). Intraperitoneal injection is preferred because it is a synchronized and effective infection route that shortens the time to produce disease symptoms, decreasing the cost of trials (Cardella and Eimers, 1990; Meza et al., 2019). Vaccine protection efficacy has been found to be affected by the route of infection for furunculosis (Midtlyng, 2005) but not for Piscirickettsiosis in Atlantic salmon (18).

On the other hand, coinfection with other pathogens such as sea lice is usually not considered in the evaluation of *P. salmonis* vaccine efficacy in laboratory-controlled conditions. We consider that this overestimates the true ability of vaccines to control Piscirickettsiosis for three reasons: first, sea lice are highly prevalent in the ocean; second, the long culture times in the sea ensure that fish will be infected not once only but several times by this pathogen; third, it has been shown that sea lice can override the protective effects of vaccination (Figueroa et al 2017). We observed no clinical sign associated with *P. salmonis* in the control tank, and mortality was significantly lower in the control tanks (less than 2–%; 3 of 137 fish) than in the cohabitating plus coinfection treatment (36–40%). Because we did not observe differences in mortality or clinical signs between vaccinated and unvaccinated adult fish in the cohabitating treatment, we predict that the evaluated vaccine will not protect fish in the field.

The immune mechanisms involved in vaccine protection against *P. salmonis* are poorly understood. In this research, the vaccine was able to induce an increase of spIgs in vaccinated fish. However, this occurred before the challenge with *P. salmonis.* After the challenge, cohabiting fish showed only increases in total Igs (41 dpi) and even a decrease of spIgs against *P. salmonis* to 41 dpi, perhaps due to B cell depletion. Apparently, the vaccine is not able to activate components of acquired immunity such as specific antibodies or cytokines associated with Th1 profiles (TNFα and IFNγ) once fish face *P. salmonis* infection, perhaps because *P. salmonis* is an intracellular pathogen. This suggests that the vaccine could act as an immunostimulant for the adaptive response at early time points, but not as a vaccine that induces future specific secondary responses. It has already been reported that vaccines may induce weaker or shorter-lived immunity in fish, mainly due to the low immunogenicity of the antigens used or because they cannot modulate the antigen presentation processes effectively during the different stages of immunity (Rozas-Serri et al., 2019). Therefore, the protective mechanism that Piscirickettsia vaccines might have in the field (Happold et al., 2020) needs to be clarified.

In Chile, the Agricultural and Livestock Service of Chile (SAG) authorized *P. salmonis* vaccines that meet a minimum protection of ≥70% RPS to be marketed. However, there is little evidence of their effectiveness under field conditions (Happold et al., 2020). In this study, the minimum protection of RPS ≥70% was not reproduced either against *P. salmonis* LF-89 strain or in the EM-90 strain. Unfortunately, neither the pharmaceutical companies nor the SAG (Agricultural and Livestock Service) publicly release the efficacy studies that authorize the marketing of vaccines in Chile. This prevented us from comparing our results with the studies carried out by pharmaceutical companies. Vaccine efficacy studies must be public and must consider both the genetic heterogeneity of the host and the pathogen’s heterogeneity. In fact, we do not know if the most vulnerable groups of populations have been included when the efficacy of the Piscirickettsiosis vaccine was evaluated by the SAG as recommended by the World Organization for Animal Health (OIE) or if pathogen heterogeneity was considered.

## 5 Data Availability Statement

The raw data supporting the conclusions of this article will be made available by the authors, without undue reservation, to any qualified researcher.

## 6 Conflicts of Interest

We declare that J.A.G., L.M., and P.C. provided genetic and immunological services to different salmon companies in Chile during the execution of this experiment. G.S and C.S. were employed in Salmones Camanchaca during the execution of this research. C.F., D.T., B.M-L, and B.D. declare no competing financial interest.

## 7 Author Contributions

J.A.G., C.S., C.F., G.S. conceived and designed the study with the help of PC and B.D. C.F., G.S., C.S., and J.A.G. performed the experiments. C.F. and D.T. performed the data analysis. B.M. and L.M performed the analysis of the Immunological data. D.T. and J.A.G. wrote the paper with the help of all authors.

## 8 Funding

This research study was funded by CONICYT-Chile through the project FONDECYT N°1140772 awarded to J.A.G and P.C. Furthermore, J.A.G was supported by the Cooperative Research Program Fellowships of OECD - PCI 2015-CONICYT. C.F. was supported by PUCV and CONICYT-Chile through a Postdoctoral fellowship (Proyecto VRIEA-PUCV Postdoctorado and FONDECYT N°3170744). D.T. was supported by CONICYT-Chile through a Postdoctoral fellowship (Becas-Chile N° 74170029). B.M-L was supported by the Postdoctoral program from the National Research and Development Agency of Chile (ANID-Chile 74200139).

## 9 Acknowledgments

We wish to thank all of the staff from the salmon selective breeding program at Salmones Camanchaca, with special thanks to Darwin Muñoz, Sonia Velazquez and Sergio Navarro for their professional support and collaboration in the experimental work. We would like to thank Salmones Camanchaca for providing fish, materials and logistics to perform this study. We would like to thank the staff of Neosalmon and Aquadvise for their valuable contribution to the development of experiments.

