## Supplementary material for "Commercial vaccines do not confer protection against two genetic strains of *Piscirickettsia salmonis*, LF-89-like and EM-90-like, in Atlantic salmon"

**Supplementary table 1.** Primary polyclonal antibodies used in ELISA analysis.

| Molecule | Source | Dilution | Reference |
|---|---|---|---|
| anti-Igs | Mouse | 1:1,000 | Supplementary Figure 1 |
| anti-TNFα | Mouse | 1:500 | Sahlmann *et al.* (44) |
| anti-IFNγ | Mouse | 1:500 | Sahlmann *et al.* (44) |

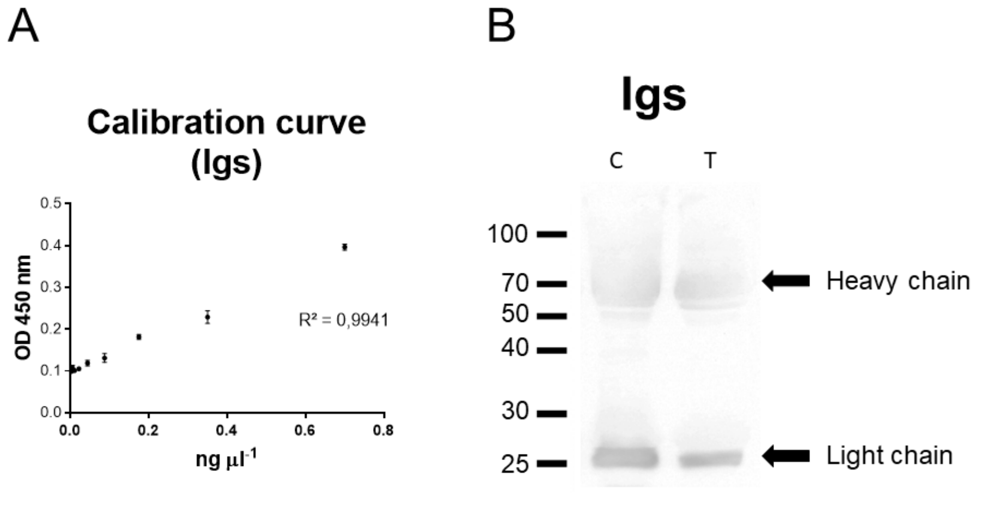


**Supplementary Figure 1.** Validation of antibodies against total serum immunoglobulins (Igs) of *Salmo salar*. A: Indirect ELISA calibration curve between total serum Igs concentration of *S. salar* (ng µL-1) and optical density at 450 nm. B: Western blot. Antibodies were produced in mice using total serum Igs from Atlantic salmon as antigen. The antigen was obtained by the caprylic acid technique for immunoglobulin purification (Fishman and Berg, 2018. DOI: 10.1101 / pdb.prot099127)

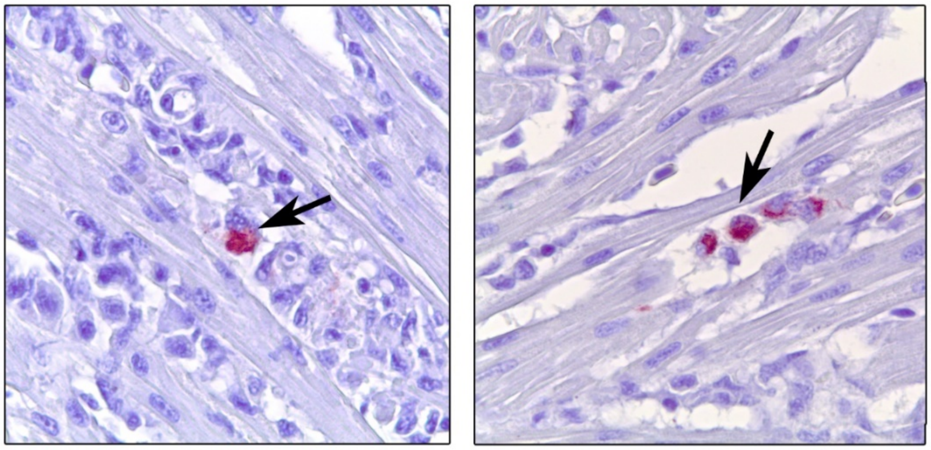

**Supplementary Figure 2.** Presence of *Piscine orthoreovirus* (black arrows) in heart samples of Atlantic salmon from the first trial at day 41 post infection with LF-89-like isolate of *P. salmonis*. The virus was detected in 7 out of 17 fish analyzed by immunohistochemistry—magnification 63X.
